# CRISPR activation of *PIKFYVE* as potential therapy for *FIG4* deficiency

**DOI:** 10.64898/2026.04.24.718784

**Authors:** Q Doctrove, GM Lenk, VH LiPuma, MH Meisler

## Abstract

*FIG4* deficiency is the cause of Charcot Marie Tooth type 4J, a neurological disorder characterized by enlarged lysosomes. Our CRISPR activation genome wide screen found that upregulation of *PIKFYVE* rescued the enlarged lysosome phenotype in cultured cells. To assess *PIKFYVE* upregulation treatment in vivo, we generated *Fig4* deficient mice with CRISPR activation of *Pikfyve* in neurons. *Pikfyve* was increased 2 fold in whole brain of CRISPR activated mice. *Pikfyve* upregulation did not extend the 3 week survival of *Fig4* deficient mice. Vacuolization of brain was not rescued. The data demonstrates that a 2 fold increase of *Pikfyve* is not sufficient to treat *Fig4* deficiency. Further testing will be required to determine if a higher increase of *Pikfyve* can ameliorate the effects of *FIG4* deficiency in vivo.

## Introduction

Charcot Marie Tooth type 4J (CMT4J) is a rare neurological disorder characterized by motor disfunction, bone deformities, and/or cognitive impairment (Chow et al., 2007; Boura et al., 2024; Sadjadi et al., 2024). These symptoms are caused by axonal loss and myelin degeneration that occur throughout the nervous system (Chow et al., 2007). CMT4J is caused by loss of *FIG4* and decreased generation of signaling molecule PI(3,5)P_2_. The PI(3,5)P_2_ signal is responsible for lysosomal maintenance. Without PI(3,5)P_2_, the lysosome becomes more acidic and enlarged with impaired function (Chow et al., 2007; McCartney et al., 2013).

To identify novel mammalian genes involved in *FIG4* function, the Meisler lab carried out a genome wide upregulation screen in *FIG4* KO HAP1 cells using CRISPR activation (Meisler et al., personal communication). *FIG4* KO cells were transduced with a lentivirus library containing sgRNAs that target 19,050 protein coding genes in the human genome, enabling overexpression of each gene (Joung et al., 2017). To only upregulate ≤ 1 gene per cell, transduction was performed at a low multiplicity of infection. Cells were screened using a pH sensitive dye that accumulated in the enlarged lysosomal compartments. Fluorescence-activated cell sorting (FACs) separated cells with enlarged lysosomes from cells with rescued small lysosomes. Rescued cells were analyzed by next gen sequencing to determine which upregulated genes could compensate for the enlarged lysosome phenotype caused by *FIG4* knockout. This genome-wide screen identified *PIKFYVE* as the top candidate for rescuing *FIG4* KO cells. Deep sequencing of the integrated sgRNAs from the rescued, non-vacuolated population revealed that *PIKFYVE* sgRNAs were significantly enriched compared with baseline (log 2 = 5.0; FDR adjusted P = 1 × 10 −10.5).”

The results of this screen suggest that *PIKFYVE* could be a treatment for individuals with deficiency of PI(3,5)P_2_. The PIKFYVE protein is the kinase directly responsible for adding a phosphate group to PI3P, converting it into PI(3,5)P_2_. PIKFYVE activity is less efficient if it is not part of the FIG4/VAC14 complex (Lees et al., 2020). However, overexpression of *PIKFYVE* in *FIG4* null cells generated enough PI(3,5)P_2_ to rescue lysosomal osmoregulation. No relevant literature exists on the effects of *Pikfyve* overexpression in vivo. The goal of our experiments was to overexpress *Pikfyve* in *Fig4* deficient mice and to evaluate the in vivo effects.

Direct administration of the *Pikfyve* mRNA is not achievable because it is too large to inject via current viral vectors. The AAV capacity is 4.8 kb and the size of the *Pikfyve* gene is 6.2 kb. We therefore investigated CRISPR activation (CRISPRa) of *Pikfyve*. CRISPRa upregulates *Pikfyve* with two components: a dead Cas9 enzyme (dCas9) fused to a transcription factor and a single guide RNA (sgRNA) that binds to and guides the CRISPRa complex to the Pikfyve gene.

Expression of *FIG4* specifically in neurons rescued *Fig4* null mice (Ferguson et al., 2012; Winters et al., 2011). We therefore tested overexpression of *Pikfyve* in neurons. AAV9 was chosen as the delivery system for the sgRNA expression because of its neuronal tropism. *Pikfyve* sgRNAs were tested in cells. One sgRNA was administered via intracerebroventricular (ICV) injection. Survival and brain histology were used to evaluate the therapeutic effectiveness of Pikfyve overexpression in *Fig4* deficient mice.

## Materials and Methods

### *Pikfyve* sgRNA sequence

sgRNAs of 20 base pairs in length, complementary to genomic sequences within 300 bp of the *Pikfyve* transcription start site, were designed with the CRISPick program (Doench et al., 2016; Sanson et al., 2018). The program used the mouse GRCm38 genome to determine the transcription start site of *Pikfyve*. The PAM site was “NGG” for the S. Pyogenes Cas9 variant and on target scoring followed the Hsu et al., 2013 rule set. The program Cas-OFFinder was used to evaluate potential off-targeting (Bae et al., 2014). Four of fourteen designed sgRNAs had a single off-target sequence with 2 base pair mismatches.

### Generation of *Pikfyve* sgRNA plasmids for in vitro evaluation

Enhanced green fluorescent protein (EGFP) was PCR amplified from the Clontech pEGFP-N2 plasmid (catalog #6081-1) and cloned into an sgRNA-MS2 backbone plasmid (Addgene #61424) according to Sambrook et al., 2001. Each designed *Pikfyve* sgRNA was inserted into an sgRNA-EGFP backbone according to the protocol of Konermann et al., 2015.

### Cell Transfection

Neuro-2a (N2A) cells (CCL-131) were obtained from The American Type Culture Collection. N2A cells were cultured in DMEM medium (Invitrogen 10569010) with 10% FBS (Sigma-Aldrich F0926) and 1x anti-anti (Gibco 15240-062). At 50-70% confluency in a 6 well plate, cells were transfected following the protocol for X-tremeGENE HP DNA Transfection reagent (Sigma-Aldrich 6366244001). Briefly, 750 ng DNA, 200 ul Opti-MEM (Gibco 31985062), and 4 ul of X-tremeGENE HP DNA transfection reagent were incubated at room temperature for 20 minutes then added to N2A cells. Two biological replicates were co-transfected with 0.5 ug of dCas9-VP64 plasmid (Addgene #61425) and 0.25 ug of *Pikfyve* sgRNA plasmid. Cells were transfected for 48 hours and RNA was extracted using TRIzol reagent (Invitrogen, 15596026)

### Quantitation of *Pikfyve* transcripts in transfected N2A cells

RNA from whole brain or transfected N2A cells was extracted with TRIzol reagent (Invitrogen 15596026) and a Direct-zol RNA MiniPrep Plus kit (Zymo R2072). Total RNA was reverse transcribed using NEB Lunascript RT Supermix Kit (E3010) according to the manufacturer’s protocol. Expression of *Pikfyve* and *Tata binding protein* was quantified by Taqman gene expression. Reactions of twenty ul contained 10 ul Taqman Gene Expression Master Mix (Invitrogen 4369016), 1 ul Taqman Gene Expression Assays for *Pikfyve* (FAM-Mm01257047_m1), and 1 ul Tata binding protein (VIC-Mm01277042_m1). Tata binding protein was used as an internal control to standardize *Pikfyve* expression. FAM and VIC fluorescence were measured with a BIO-RAD CFX96 C1000 TOUCH Real-Time PCR System. Gene expression was quantified by comparing non-transfected to sgRNA transfected samples using the ΔΔCT method (Livak and Schmittgen, 2001).

### Mice

The spontaneous *Fig4* null mutation, pale tremor (Fig4^plt^, JAX 017800), was identified in our laboratory (Chow et al., 2007). The *Fig4* mutation causes tremors, motor dysfunction, a pale coat color, and death at three weeks postnatal. This mouse has been backcrossed for more than 20 generations to strain C57BL/6J (JAX 000664).

Conditional *Fig4* knockout mice (Fig4^flox^, JAX 039611) with loxP sites flanking exon 4 of the *Fig4* gene were generated in our lab by targeted germline mutation in embryonic stem cells (Ferguson et al., 2012).

Mice that conditionally express dCas9-VPR (dCAM) were generously provided by Dr. Wolfgang Wurst, Ludwig Maximilian University of Munich (Giehrl-Schwab et al., 2022). In this line, the dead Cas9 protein is fused to the transcriptional activators VP64, p65, and Rta. A Lox-Stop-Lox sequence is located upstream of dCas9-VPR. In the presence of Cre-recombinase the Lox-Stop-Lox sequence is deleted and dCAM expression is activated (Fig 4A).

Nestin-Cre mice (Jax line 003771) express Cre-recombinase in neuronal progenitor cells and their derived cells in the central and peripheral nervous system. This Cre recombinase becomes active at embryonic day 9-14 (Mouse Genome Informatics database, Baldarelli et al., 2024).

Mice were crossed to generate animals heterozygous for two mutant *Fig4* alleles (null/floxed), heterozygous for Nestin Cre (Nestin/+), and heterozygous for the dCas9-VPR (dCAM/+) (Fig. 4). These mice express dCAS9-VPR and lack *Fig4* in the central and peripheral nervous system. For genotyping primers see table 3-1.

### Intracerebroventricular (ICV) injection of AAV9-sgRNA

P0 to P2 mice were cryoanesthetized and injected with 2 ul of 4.4E+13 vg/ml sgRNA AAV9 in the right ventricle. Free hand injection was performed using a 5 ul syringe (Hamilton 7634-01) and a 33-gauge needle (Hamilton 7803-05). Coordinates for the targeted injection were 0.25 mm lateral from the sagittal suture, 0.5 mm anterior to the lambdoid suture, and 2 mm deep.

### Brain Histology

After in vivo expression of AAV9 sgRNA, brain vacuolization was examined by H&E staining (Histoserv Bethesda, MD).

mKATE fluorescence was examined in brain tissue. Tissue was incubated overnight in paraformaldehyde at 4 degrees Celsius followed by incubation in 30% sucrose for another 24 hours. Brain was frozen in O.C.T (ThermoFisher 4585) and 12-micron sections were prepared with a Leica CM1950 Cryostat. Tissue slices were mounted on slides using Prolong Gold antifade reagent with DAPI (Invitrogen P36931). mKATE fluorescent images were taken using Nikon Ti2 microscope at 20x and 40x.

## Results

### Identification of sgRNAs that upregulate *Pikfyve* in cultured cells

The CRISPick program from the Broad Institute (Doench et al., 2016; Sanson et al., 2018) was used to design sgRNAs targeting the region from 0 to -300 base pairs upstream of the *Pikfyve* transcription start site (Fig. 1A). Each sgRNA was ligated into a plasmid driven from a U6 promoter as described by Konermann et al., 2015. Each sgRNA plasmid was co-transfected with a plasmid expressing dCas9-VP64 in mouse neuroblastoma Neuro-2A (N2A) cells. Endogenous *Pikfyve* expression was quantified by qRT-PCR (Fig. 1B). sgRNA 9, located from - 141 bp to -161 bp upstream of the transcriptional start site, had a 2.5 fold increase of *Pikfyve* expression in comparison with no sgRNA (Fig. 1B, asterisk). This sgRNA was selected for *in vivo* testing.

**Figure 1.**
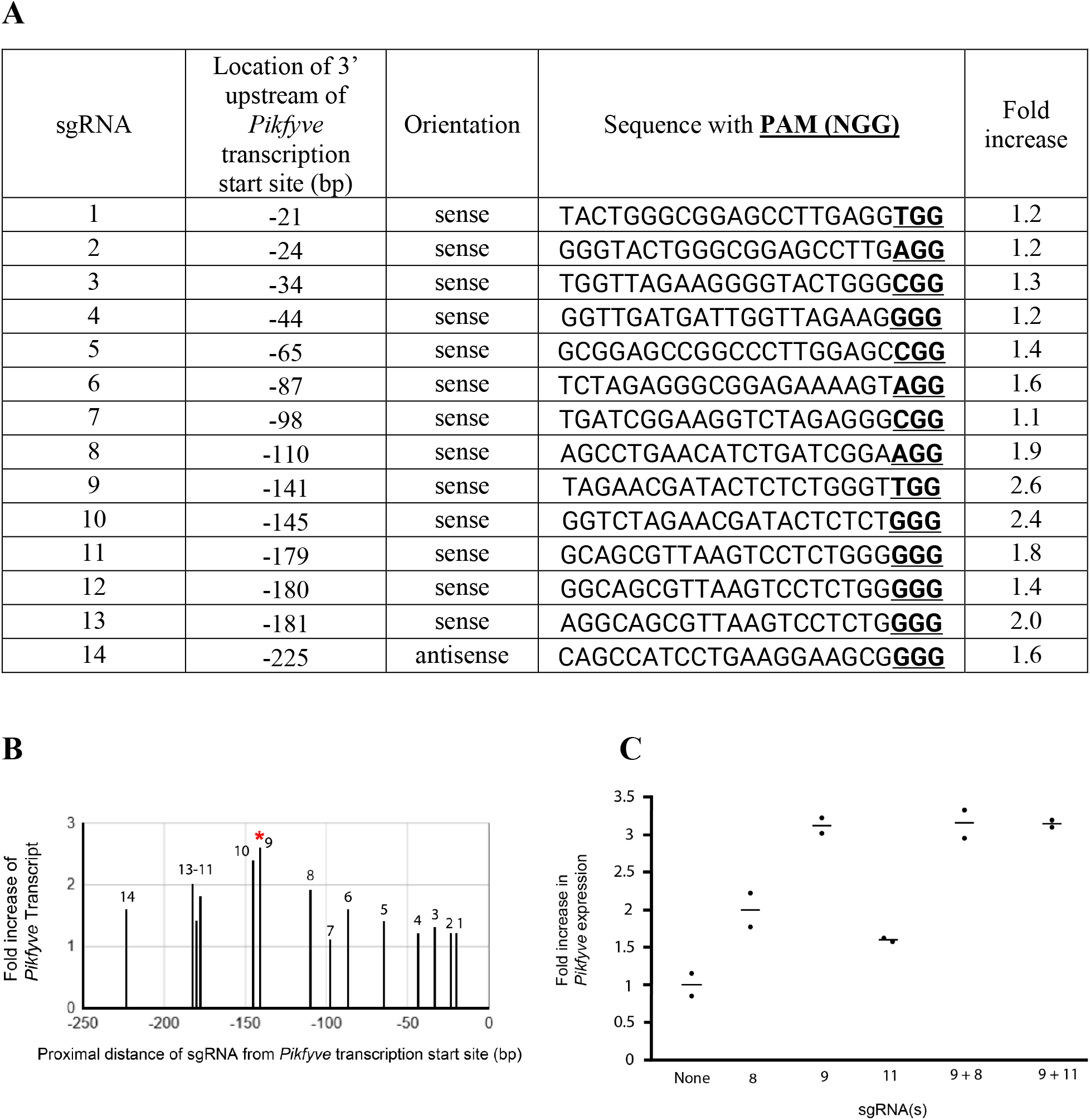
CRISPRa upregulation of *Pikfyve* in N2A cells. (A) Fourteen sgRNA sequences were designed up to - 300 base pairs upstream of the *Pikfyve* transcriptional start site. (B) Upregulation of *Pikfyve* in N2A cells was evaluated using *Pikfyve* Taqman assay and normalized with*Tata binding protein* Taqman assay. The asterisk labels the highest expressing sgRNA used in later experiments. (C) Combinations of sgRNAs

We attempted to further increase *Pikfyve* expression by combining sgRNAs. Combining sgRNA 9 with sgRNA 8 or sgRNA 11 did not further increase the expression of *Pikfyve* (Fig.1C). Co-transfection of SAM components did not significantly increase expression (not shown).

### *Fig4* knockout mouse with CRISPR activation of *Pikfyve* in neurons

To determine if *Pikfyve* upregulation can rescue *Fig4* deficiency *in vivo*, we generated mice that are *Fig4* null in neurons and express CRISPRa components in neurons. To accomplish this, we genetically combined the *Fig4* null (Chow et al., 2007), *Fig4* flox (Ferguson et al., 2012), dCas9 activator (dCAM)(Giehrl-Schwab et al., 2022), and Nestin-Cre mice (Baldarelli et al., 2024).

*Fig4* knock out mice exhibit phenotypes similar to *Fig4* deficient patients. Mouse phenotypes include reduced body weight, tremor, vacuolization of neurons in the brain and dorsal root ganglia, and a reduced lifespan of about 6 weeks (Chow et al., 2007). *Fig4* flox mice were included in the cross to localize complete knock out *Fig4* to neurons in the presence of Nestin-Cre.

dCAM mice express the CRISPR activation components inserted at the Rosa26 locus (Fig. 2A), in the presence of Cre recombinase. Nestin-Cre successfully removed the flox stop flox sequence, activating dCas9 VPR in brain lysate (Fig. 3). To determine the dCAM genotype, dCAM primers amplified a 454 bp fragment of the dCas9 sequence, while wildtype primers amplified a 300 bp fragment of the WT Rosa26 locus (Fig. 2; Table 1) Activated dCAM mice were designed to express dCas9-VPR (Chavez et al., 2015) and synergistic activation mediator (SAM) (Konermann et al., 2015). Long read sequencing captured the dCAM sequence, including the CAG promoter and 1.2 kb of the dCas9, revealing that the SAM components were absent (Fig. 3). The dCas9 sequence remained as intended (Fig. 3C), thus our experiment proceeded as planned only without the SAM activation.

**Table 1.**
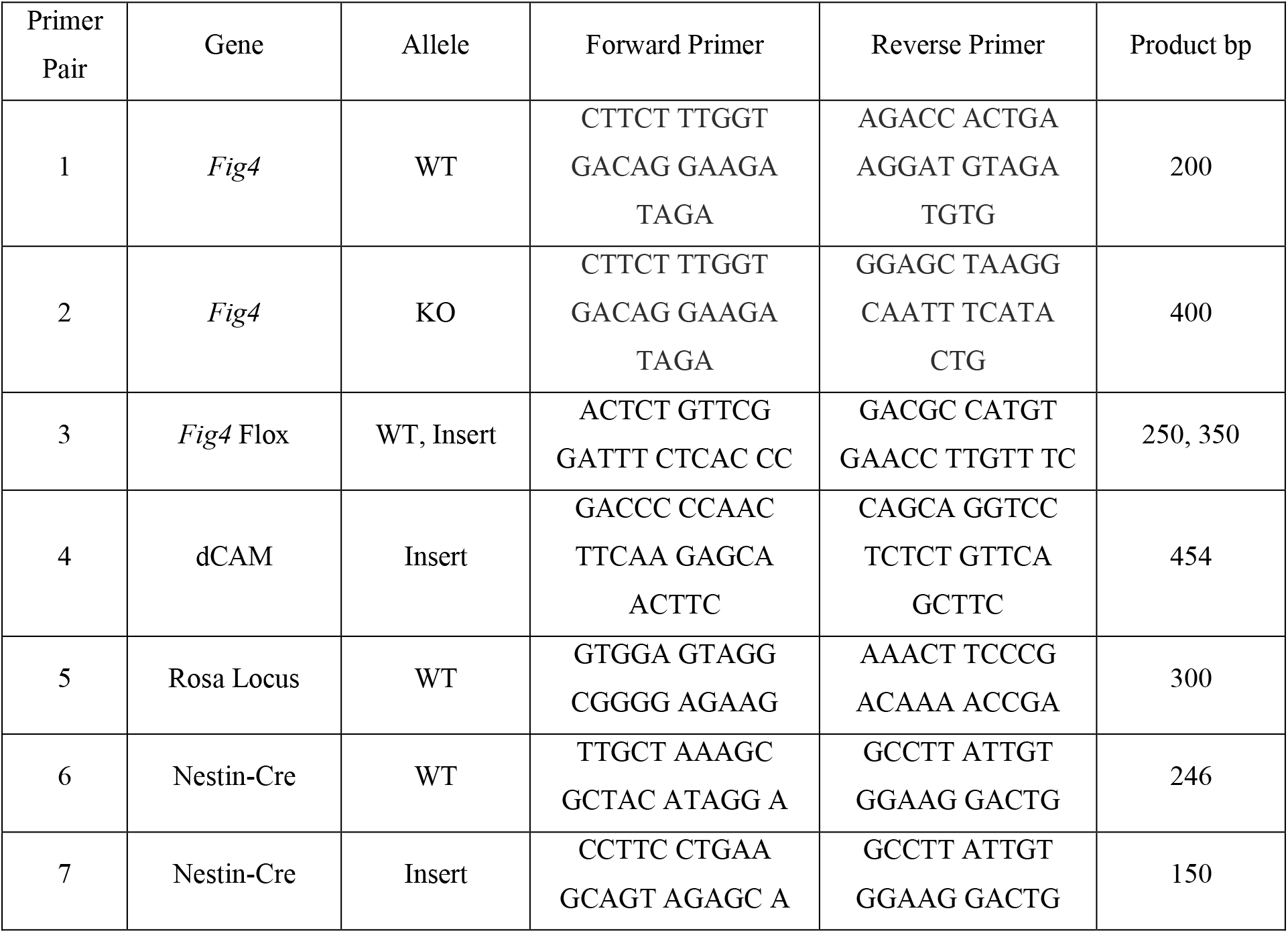
Genotyping primers. Sequences are continuous.

**Figure 2.**
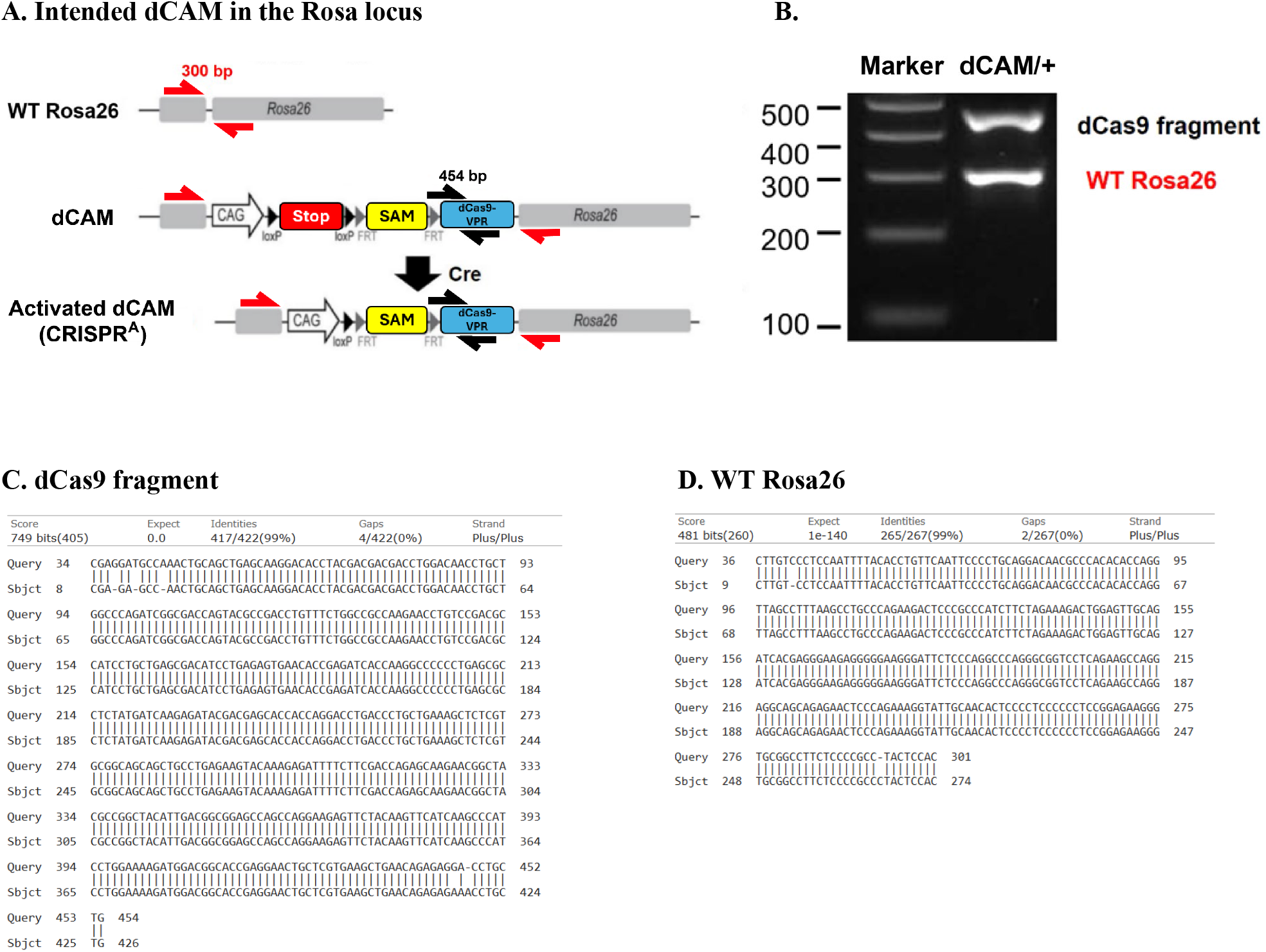
Proposed dCAM transgene in the mouse Rosa locus. (A) Prior to Cre activation, the dCAM allele does not express the CRISPRa components due to the presence of the stop cassette. Crossing the dCAM mouse with a Cre-recombinase expressing animal generates the CRISPRa expressing animal. SAM contains the transcriptional activator fusion protein MS2-HSF1-P65. Red arrows: WT Rosa26 genotyping primers. Black arrows: dCas9 genotyping primers. Adapted from Giehrl-Schwab et al., 2022. (B) dCAM genotyping of a heterozygous dCAM mouse. (C) dCas9 and (D) WT PCR validated using sanger sequencing. Query is dCas9 or WT Rosa26 sequence, Sbjct is sanger sequencing of PCR products.

**Figure 3.**
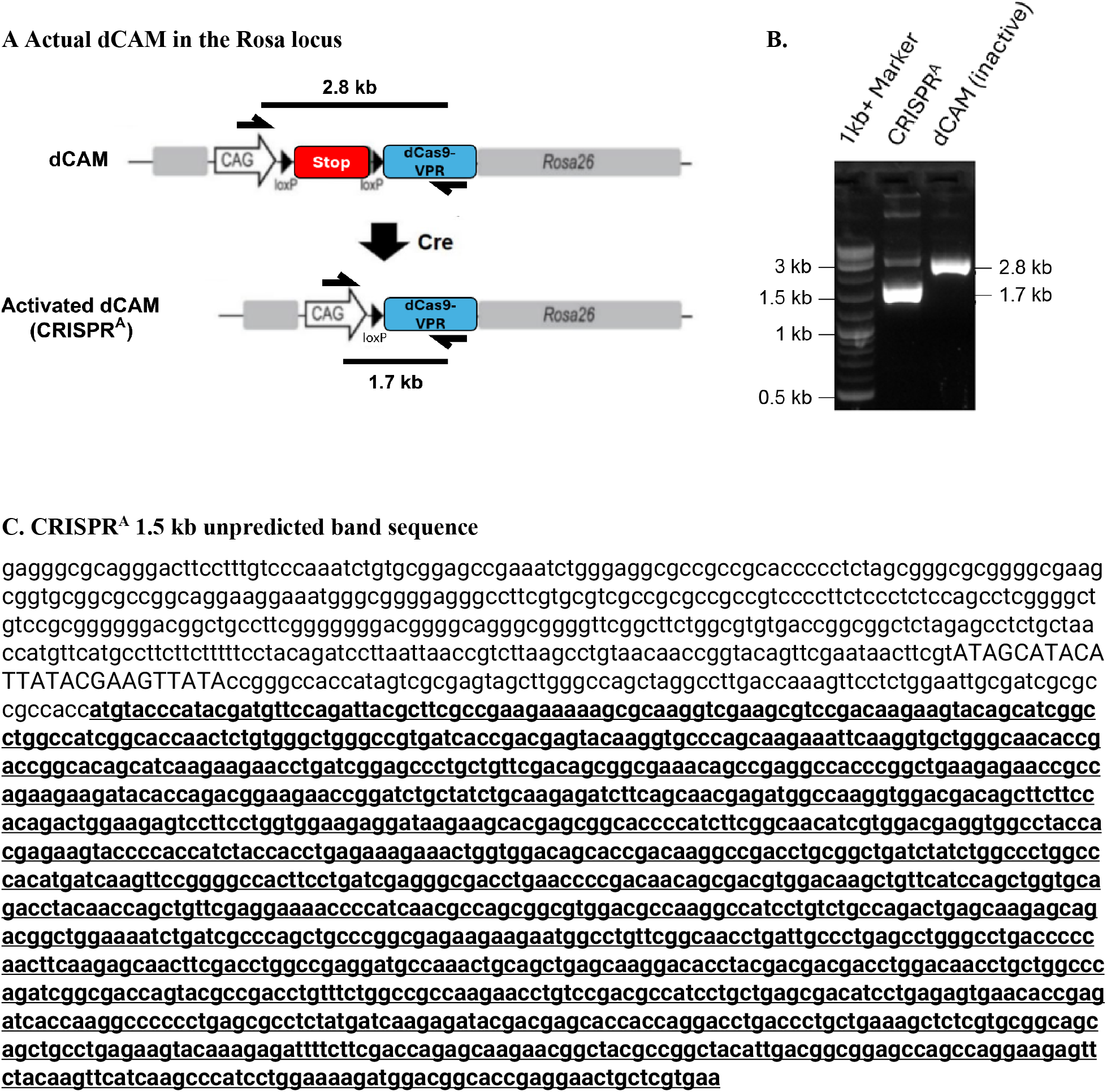
The SAM transcription factor components are missing from dCAM animals. (A) Primers were designed to genotype inactive dCAM or CRISPR^A^. Black arrows: CRISPR^A^ primers. Inactive dCAM PCR product was predicted to be 2.8 kb and CRISPR^A^ PCR product was predicted to be 1.7 kb. (B) The genotyping assay of a CRISPR^A^ (dCAM/+, Nes/+) mouse. (C) The 1.7 kb CRISPR^A^ band was subjected to long read sequencing. SAM transcription factor sequence is missing but dCas9 sequence remains. Uppercase, remainder of loxp sequence.Underlined and bold, captured dCas9 coding region.

Five crosses were performed to generate the experimental and control mice (Fig. 4).Homozygous *Fig4* null mice cannot be used in crosses as they die before sexual maturity, thus heterozygous *Fig4* null were used for breeding. We generated dCAM homozygous mice that were heterozygous for the flox allele of *Fig4* (Cross 1a and 1b). Simultaneously we generated Nestin-cre homozygous mice that were heterozygous for *Fig4* null (cross 2a and 2b). Crossing the offspring from 1b and 2b together generated the experimental group (dCAM/+, Nes-Cre/+, *Fig4* null/floxed) and controls (dCAM/+, Nes-Cre/+, *Fig4* +/+). Both experimental and control groups express dCas9 VPR in neurons as both groups express one allele of Nestin Cre and one allele of dCAM. The final yield of Fig4^flox^/Fig4^-^ mice from cross 3 was 15/57, consistent with Mendelian prediction of 1/4 (p=0.71, Fisher’s exact test).

**Figure 4.**
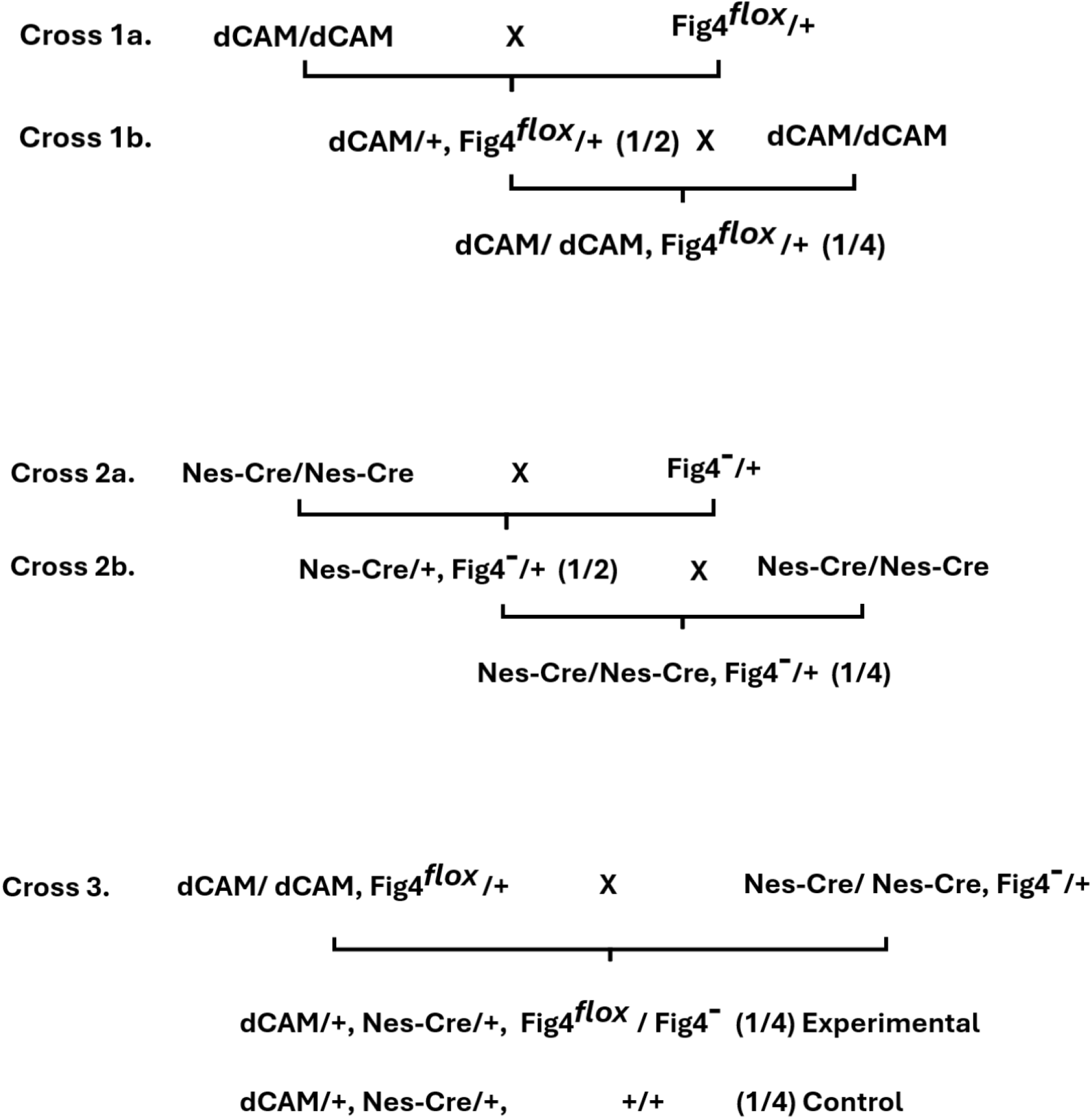
Generation of dCAM activated mice using Nestin-Cre with and without *Fig4* expression. The final yield of Fig4^flox^/Fig4^-^ mice from cross 3 was 15/57, consistent with Mendelian prediction of 1/4 (p=0.71, Fisher’s exact test)

### Administration of sgRNA to *Fig4* mutant mice

An AAV9 construct for administering the *Pikfyve* sgRNA was generated by the University of Michigan Vector Core (Fig. 5). AAV9 was chosen for its tropism for neurons (Issa et al., 2023). In this construct sgRNA 9 is expressed from a U6 promoter. The red fluorescent marker mKATE is expressed from a ubiquitous CAG promoter. Experimental and control animals from cross 3 (Fig. 4) were injected with 2 ul of AAV9 (4.4×10^13^ vg/ml) via ICV injection. Activation of dCAM by Nestin-Cre was confirmed by RT-PCR using RNA from mouse brain lysate (Fig. 6A). Successful injection and expression of AAV9 was confirmed by the presence of mKATE-NLS fluorescence in all brain slices evaluated (Fig. 6B). Expression of endogenous *Pikfyve* in whole brain at P14 was quantified by qRT-PCR (Fig. 7). sgRNA-injected mice exhibited a 2 fold increase of *Pikfyve* expression compared with noninjected controls.

**Figure 5.**
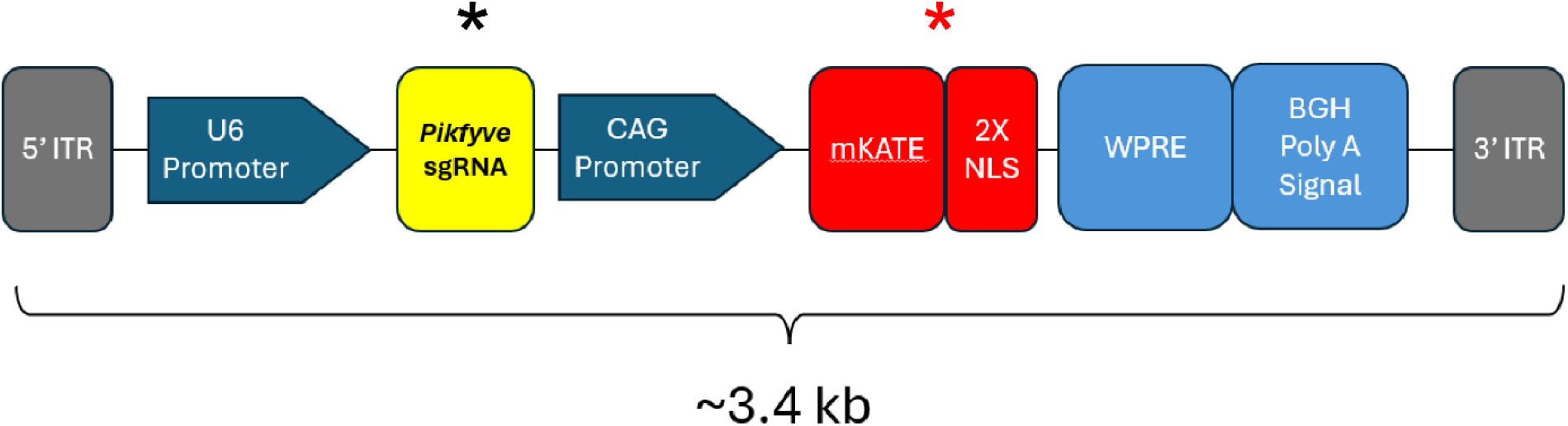
Structure of AAV9 for delivery of *Pikfyve* sgRNA and mKATE marker. *Pikfyve* sgRNA 9 (black asterisk) is driven from a U6 promoter. Red fluorescent marker protein mKATE (red asterisk) is fused to nuclear localization signals and driven from a ubiquitous CAG promoter.

**Figure 6.**
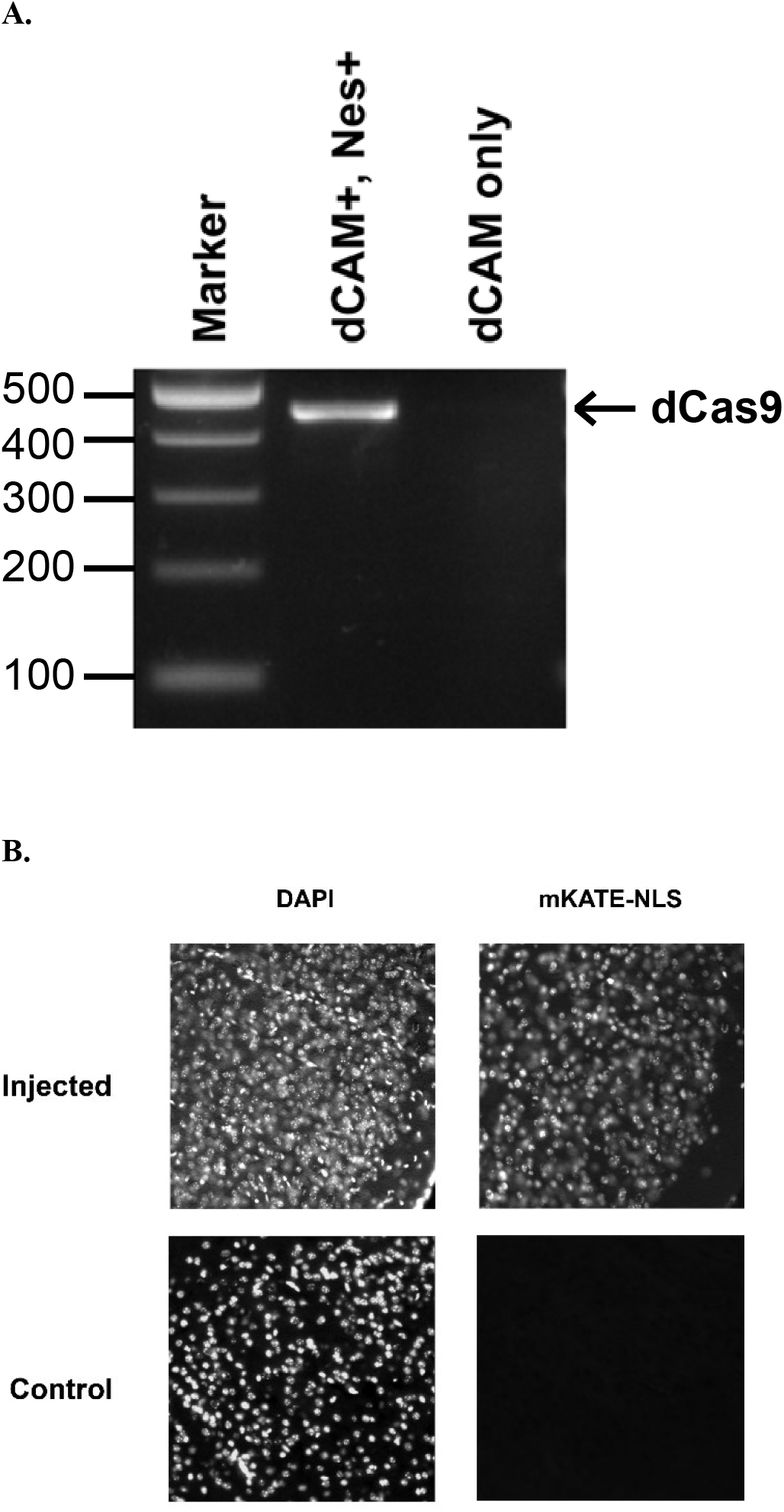
dCas9 and mKATE (a marker of Pikfyve sgRNA AAV9) are expressed in brain of injected animals prior to lethality. (A) The RT-PCR product of dCas9 is present in mouse brain lysate when *Nestin* (Nes+) is expressed at P14. (B) Mice that were injected with sgRNA AAV9 express the mKATE marker (Fig. 5) in cortex.

**Figure 7.**
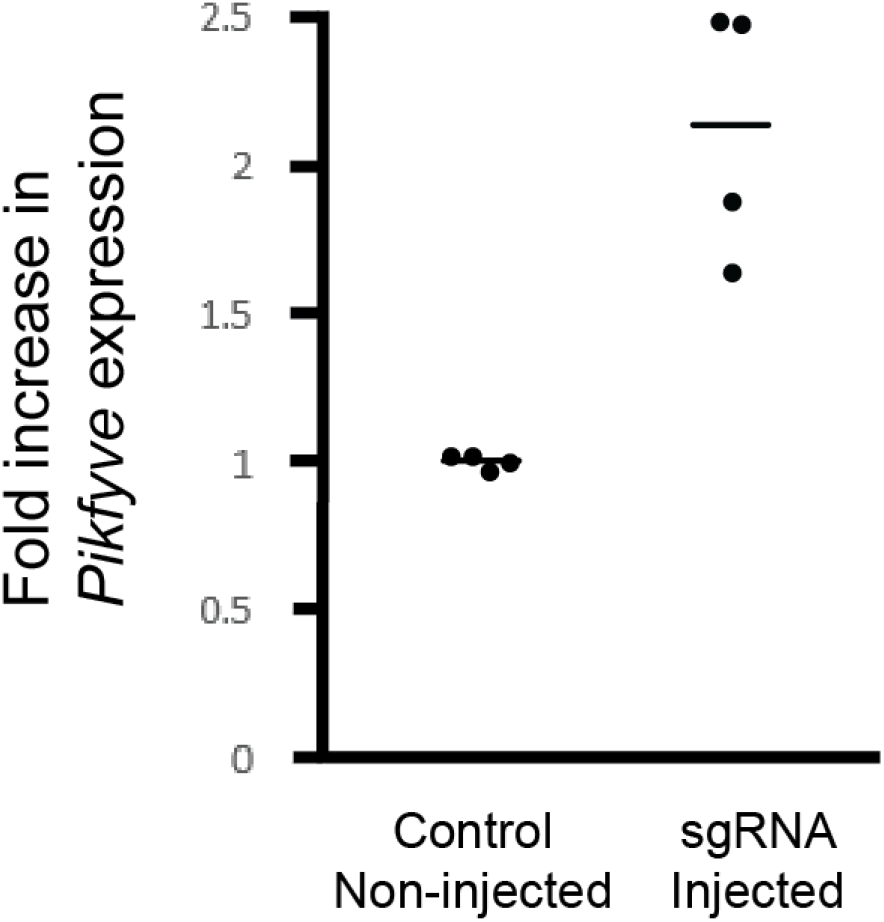
*Pikfyve* mRNA expression in sgRNA injected animals. qRT-PCR of whole brain mRNA was evaluated using *Pikfyve* Taqman assay and normalized to *Tata binding protein* Taqman assay. (p<0.001, Unpaired t test)

### Effects of CRISPRa treatment

The survival of sgRNA-injected mice was not extended beyond the three-week survival of non-injected control mice (Fig. 8). Enlarged vacuole phenotypes in brain were also not rescued in sgRNA injected animals (Fig. 9). Thus the 2 fold increase of *Pikfyve* expression did not rescue the *Fig4* deficient mice. We attempted, unsuccessfully, further increase the expression of *Pikfyve* by expression of additional transcription factors or by combining multiple sgRNAs. It remains possible that higher expression of *Pikfyve* might rescue *FIG4* deficiency, as originally proposed.

**Figure 8.**
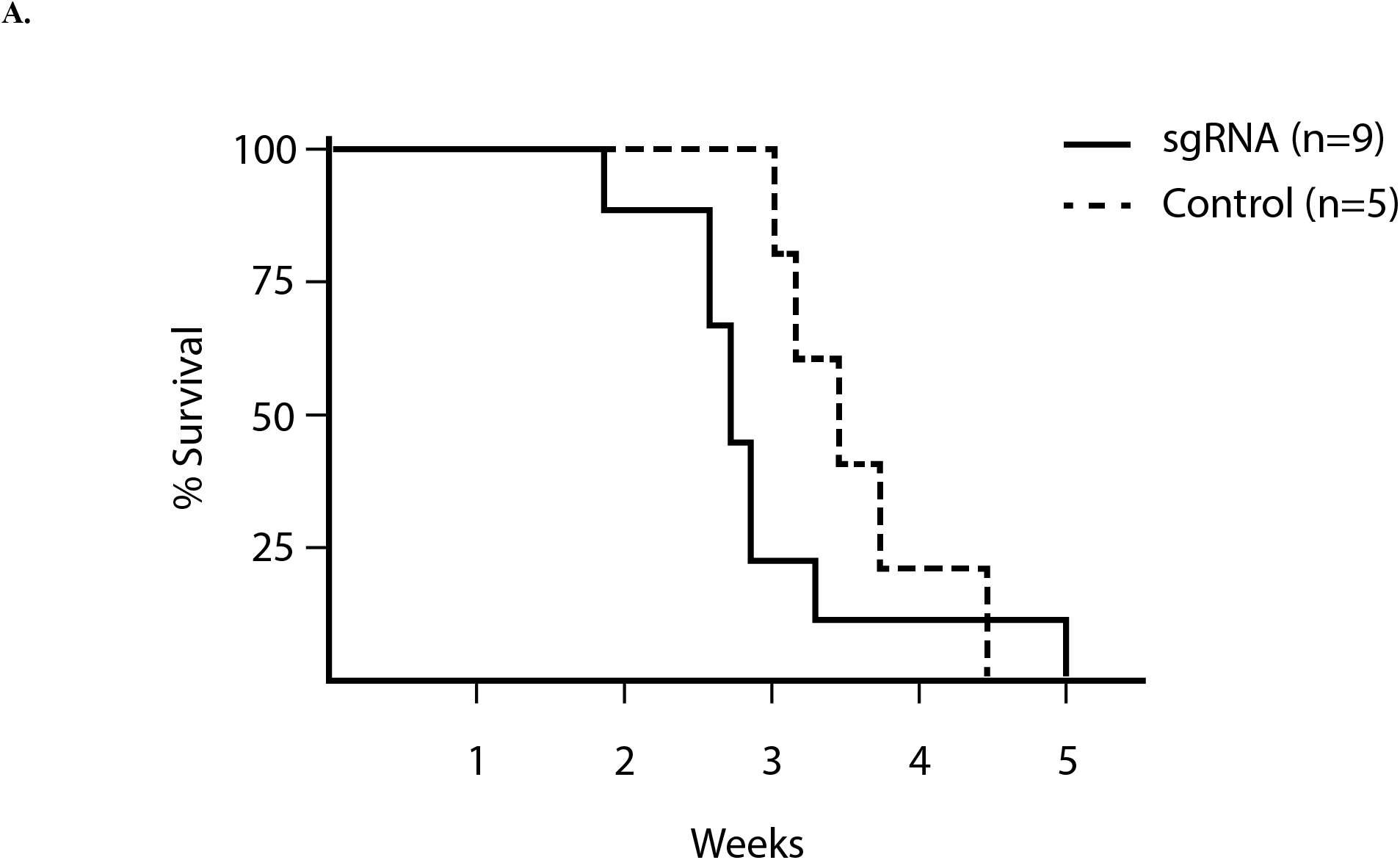
Treatment of sgRNA does not rescue survival of *Fig4* deficient mice. The survival of sgRNA-AAV treated Fig4^flox^/Fig4^-^ mice (solid line) did not differ significantly from non-injected control mice (dashed line) (Log rank test, *p* = 0.28)

**Figure 9.**
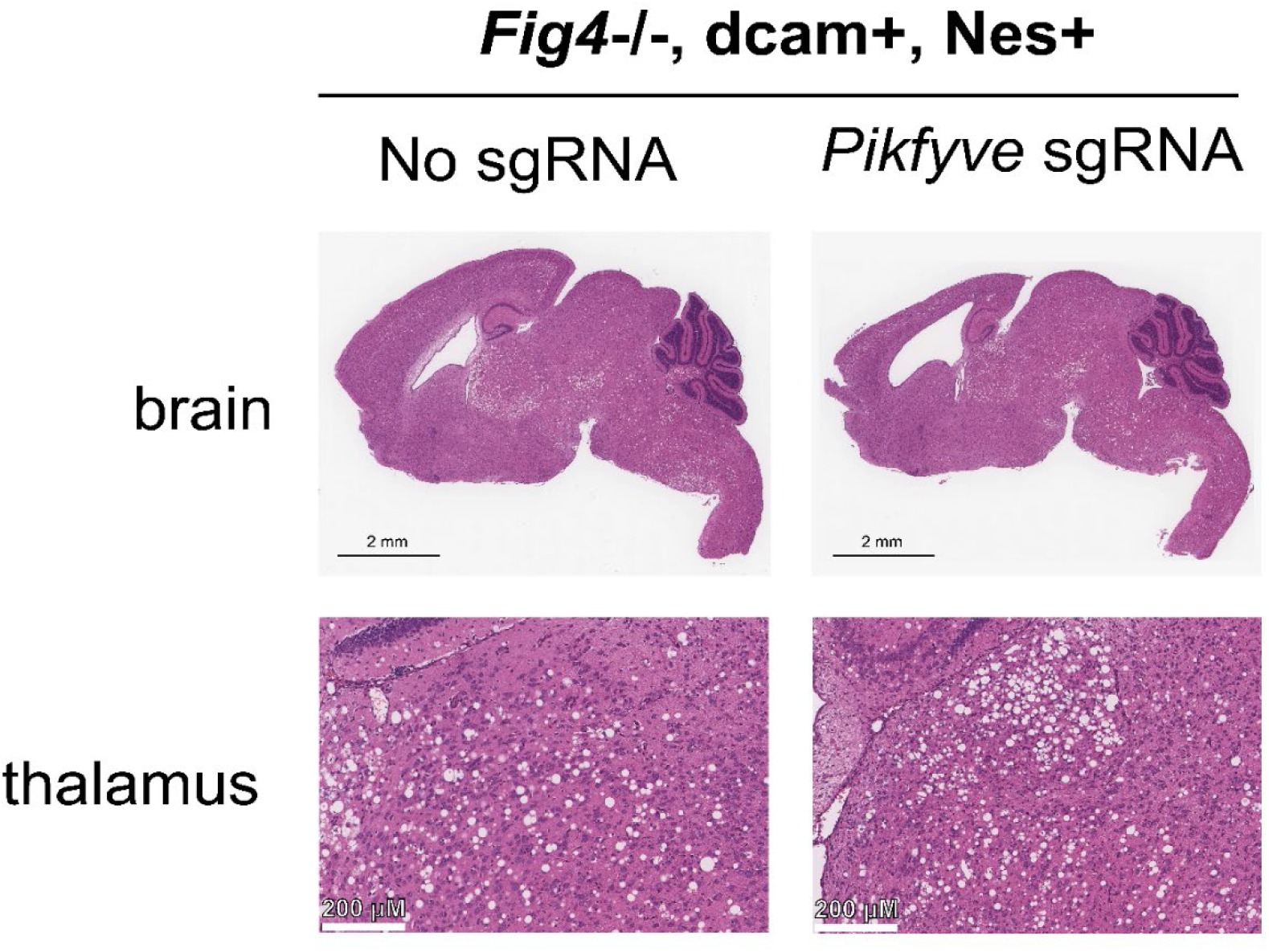
Treatment of sgRNA does not rescue vacuolization of *Fig4* deficient mice. Brains from P14 mice were stained with hematoxylin and eosin.

## Discussion

CRISPR activation offers a new system for studying disease and developing treatments by overcoming the current packaging size limit for large gene expression (Perez-Pinera et al., 2013, Chavez et al., 2015, Konermann et al., 2015). Recent studies demonstrate successful in vivo treatment of neurological disorders using CRISPR activation, including epilepsy (Colasante et al., 2019; Tamura et al., 2025), Parkinson’s disease (Giehrl-Schwab et al., 2022), and Fragile X syndrome (Geng et al., 2025). Our genome wide screen demonstrated that upregulation of *PIKFYVE* using CRISPR activation rescues the enlarged lysosome phenotype in *FIG4* null cells. This indicates PIKFYVE retains the ability to generate some signaling molecule PI(3,5)P_2_ without FIG4 stabilization. In this study, we investigated whether CRISPR activation of *Pikfyve* could compensate for loss of *Fig4* in vivo using the CMT4J mouse.

To upregulate *Pikfyve* in our *Fig4* deficient mice, sgRNAs were designed and tested in mouse neuroblastoma cells (N2A). The greatest increase in expression was 2.5 fold. sgRNA was delivery by AAV9 intracerebroventricular injection, a method previously shown to transduce affected neuronal populations and rescue *Fig4* deficiency through gene replacement (Presa et al., 2021). We generated mice expressing conditional *Fig4* knockout alleles with a Cre activated CRISPR activation system. Nestin-Cre was used to knockout *Fig4* and drive expression of the dCas9-VPR activator in neuronal progenitors and their descendants. Long read sequencing revealed that the dCAM allele used in this study lacked the intended SAM components, leaving only the dCas9-VPR transcriptional activator. The absence of SAM likely reduced the strength of CRISPR activation. Despite this limitation, AAV9 delivery of sgRNA resulted in a 2 fold increase in *Pikfyve* expression in the brain, confirming that the CRISPR activation system was functional in neurons.

The increase in *Pikfyve* expression did not improve neuropathological phenotypes in *Fig4* knockout mice. Treated animals did not have an extended lifespan or reduction in neuronal vacuolization. A 2 fold increase in *Pikfyve* expression was insufficient to compensate for the loss of *Fig4* function in this model. A substantially larger increase in *Pikfyve* activity may be required to restore PI(3,5)P_2_homeostasis in the absence of *Fig4*. Alternatively, *Fig4* may have functions beyond its role in the *Pikfyve* stabilization, which cannot be compensated for by increasing *Pikfyve* levels alone.

Several technical factors may have limited the effectiveness of *Pikfyve* activation in vivo.The absence of SAM components in the dCAM mice likely reduced the magnitude of transcriptional activation achievable with the CRISPR system. *Pikfyve* expression was measured in whole brain lysates, which could underestimate the degree of activation in specific neuronal subpopulations if transduction efficiency varied across regions or cell types.

Future studies could address these limitations by employing stronger transcriptional activation systems, including full CRISPR SAM or alternative activator techniques, to achieve higher levels of *Pikfyve* expression. Targeting multiple regulatory regions of the *Pikfyve* locus or using improved sgRNA designs may also enhance transcriptional activation.

In summary, this study demonstrates that CRISPR activation can be used to increase *Pikfyve* expression in neurons in vivo. However, the modest upregulation achieved here was insufficient to rescue the pathological phenotypes of *Fig4* deficiency. These findings suggest that stronger activation of *Pikfyve* may be required to ameliorate disease. The system I have developed could be used to test higher levels of *Pikfyve* upregulation, test other genes from our overexpression screen that may rescue *Fig4* deficiency (such as FAM241, SLC12A9, and SLC2A6), and evaluate disease phenotypes in other mouse strains (such as Vac14 null mice (Jin et al., 2008)).

